# Age-associated transcriptomic and epigenetic alterations in mouse hippocampus

**DOI:** 10.1101/2024.09.05.611100

**Authors:** Merve Bilgic, Yukiko Gotoh, Yusuke Kishi

## Abstract

Aging represents a major risk for human neurodegenerative disorders, such as dementia and Alzheimer’s disease, and is associated with a functional decline in neurons and impaired synaptic plasticity, leading to a gradual decline in memory. Previous research has identified molecular and functional changes associated with aging through transcriptomic studies and neuronal excitability measurements, while the role of chromatin-level regulation in vulnerability to aging-related diseases is not well understood. Moreover, the causal relationship between molecular alterations and aging-associated decline in functions of different cell types remains poorly understood. Here, we systematically characterized gene regulatory networks in a cell type–specific manner in the aging mouse hippocampus, a central brain region involved in learning and memory formation, by simultaneously profiling gene expression and chromatin accessibility at a single nuclei level. The analysis of multiome (RNA and ATAC) sequencing recapitulated the diversity of glial and neuronal cell types in the hippocampus, and allowed revealing transcriptomic and chromatin accessibility level changes in different cell types, among which oligodendrocytes and dentate gyrus (DG) neurons exhibited the most drastic changes. We found that aging-dependent chromatin-level changes were more pronounced than transcriptomic changes for genes related to synaptic plasticity among neurons. Our data suggest that BACH2, a candidate transcription factor in the aging-mediated functional decline of DG neurons, potentially regulates genes associated with synaptic plasticity, cell death, and inflammation during aging.

## INTRODUCTION

The aging brain undergoes a series of transformations involving the hippocampus that have profound implications for memory and cognitive function. The hippocampus plays a pivotal role in memory formation and retrieval, and its functional decline with aging has implications in aging-related disorders including Alzheimer’s disease (AD) due to impaired synaptic plasticity and intrinsic excitability (Nicholson *et al*, 2004; Lu *et al*, 2004; Oh *et al*, 2010). Understanding how aging-associated changes occur in the hippocampus has been important in the field of neuroscience and medical sciences.

Previous research has illuminated molecular and cellular hallmarks of aging, such as transcriptomic and epigenetic alterations, cellular senescence, and inflammation in the mammalian brain (Guo *et al*, 2022; Wang *et al*, 2022; Campisi *et al*, 2019; López-Otín *et al*, 2013; Hauss-Wegrzyniak *et al*, 2000; Murray & Lynch, 1998; Allen *et al*, 2023). Gene expression studies on the mammalian brain reflected aging-dependent biological processes including inflammation, oxidative stress, decreased mitochondrial function and protein processing (Lee *et al*, 2000; Blalock *et al*, 2003; Lu *et al*, 2004). Moreover, epigenetics contribute to cellular memory via the regulation of DNA, histone modifications, and chromatin structure, and its alteration during the aging process of many tissues is well known (Wang *et al*, 2022; Talens *et al*, 2012; Han & Brunet, 2012; Maybury-Lewis *et al*, 2021; Ucar *et al*, 2017; Ernest Palomer *et al*, 2016). For example, in the hippocampus, the inhibition of histone deacetylase (HDAC) enhanced hippocampus-dependent memory and synaptic plasticity (Vecsey *et al*, 2007; Guan *et al*, 2009), although other studies showed no difference (Harman & Martín, 2020; Sewal *et al*, 2015). In addition, manipulation of the DNA methyltransferase (DNMT) gene affects hippocampus-dependent brain function. Knockout of *Dnmt1* impaired the spatial memory formation of aged mice (Liu *et al*, 2011), and overexpression of *Dnmt3a2* in the hippocampus of aged mice restored the formation of fear memory and object location (Oliveira *et al*, 2012). However, the epigenetic alterations during hippocampal aging at the genome-wide level remain unknown, while alterations in some specific gene loci were reported (Palomer *et al*, 2016).

Nevertheless, aging is a gradual process, marked by cumulative effects of cellular damage that occur asynchronously across individuals, tissues and cells (Buckley *et al*, 2022). Comparative proteomic profiling of the hippocampus and occipital cortex synapses in primates demonstrated different age-dependent synaptic variance with a selective vulnerability in hippocampal synapses (Graham *et al*, 2019). Proteomic studies of the human AD brain and aged mouse brain identified different protein changes in vulnerable and resistant brain regions (Askenazi *et al*, 2023; Keele *et al*, 2023; Mol *et al*, 2022; Astillero-Lopez *et al*, 2022; Oliviero *et al*, 2022). Even within the hippocampus, previous research revealed that aging has different effects on different types of neurons in structurally and functionally distinct hippocampal regions, including the dentate gyrus (DG), Cornus Ammonis-1 (CA1), CA2, CA3, and subiculum (Flood *et al*, 1987a; Flood, 1991; Flood *et al*, 1987b; Hanks & Flood, 1991; Maziar *et al*, 2023). Together, these observations highlight the different vulnerabilities of neurons to the aging process and neurodegenerative diseases. Single-cell omics studies emerged as a powerful tool to profile aging-dependent transcriptomic and epigenetic changes in healthy and diseased brains at a single-cell level (Ogrodnik *et al*, 2021; Ximerakis *et al*, 2019; Luo *et al*, 2023; Shao *et al*, 2021; Zhang *et al*, 2022; Hajdarovic *et al*, 2022; Luo *et al*, 2022; Shi *et al*, 2018). Many studies identified the specific molecular state of microglia, immune residents in the brain, in normal aging and in AD conditions in mice and humans (Li *et al*, 2023; Xiong *et al*, 2023; Sun *et al*, 2023; Lopes *et al*, 2022). Others found a regional specificity in aging-dependent changes of gene expression in glial cells, non-neuronal cell types supporting neuronal development and function (Soreq *et al*, 2017; Astillero-Lopez *et al*, 2022). Excitatory and inhibitory neurons in the hippocampus of patients with AD exhibited region-specific transcriptomic changes (Luo *et al*, 2023; Mathys *et al*, 2023). In addition to single-cell transcriptomic analysis, single-nucleus ATAC-sequencing (snATAC-seq) allowed researchers to generate an atlas of cis-regulatory DNA elements (CREs) of different cell types in the adult mouse brain (Li *et al*, 2021; Zu *et al*, 2023; Yao *et al*, 2021). Investigating the specific changes that different cell types undergo as they age remains a critical step toward understanding their association with increased susceptibility to brain dysfunctions and improving the selection of appropriate therapeutic targets.

However, despite the advancements, how cell type–specific alteration of CREs during hippocampal aging and their contribution to aging-related vulnerabilities of different neuronal subtypes in the hippocampus remain less understood. To address these limitations, we investigated the dynamic alterations in gene expression and chromatin accessibility in the hippocampus by employing single-nucleus multiome sequencing of RNA and ATAC libraries from the same nuclei. This method allows us to obtain an improved sensitivity and proper specificity of CRE-gene expression links compared with computational integration of separately performed snRNA-seq and snATAC-seq assays (Xie *et al*, 2023).

To assess cell type–specific age-dependent changes, we performed a comprehensive analysis of differentially expressed genes (DEGs) and differentially accessible regions (DARs) by comparing young and aged cells for each cell type. Our results revealed that glial cells, including oligodendrocytes and astrocytes, exhibited more pronounced transcriptomic changes compared with neurons in general. Among neurons, DG neurons displayed the highest numbers of DEGs with aging, followed by CA1 and subiculum. However, rather than gene expression, most of the synaptic plasticity and other neuronal function–related genes were altered at the chromatin level. Our motif enrichment analyses revealed the BACH2 transcriptional repressor as a potential regulator of the aging process in DG neurons by regulating the epigenetic and transcriptomic landscape of synaptic and survival genes.

## RESULTS

### Single-nuclei profiling of transcriptome and chromatin accessibility in the aging mouse hippocampus

To understand the aging-associated changes of gene regulatory programs in different cell types of the hippocampus, we profiled the transcriptome and chromatin accessibility of hippocampal cells isolated from two samples of 7-week-old (young) and 108-week-old (aged) male mice (Figure 1A). We took advantage of single-cell multiomics to capture the cellular heterogeneity and minimize the variance of aging-related profiles across samples by obtaining both RNA and ATAC libraries from the same single nuclei. This method allowed a direct comparison between the gene expression and chromatin state of genes and minimized sample variance by simultaneous mapping of transcriptomic landscapes and chromatin accessibility within individual cells, hence enhancing the depth of our analysis. We filtered low-quality cells and performed dimension reduction and clustering using ArchR software (Methods; Granja et al., 2021). The quality control filtering retained 15,480 nuclei (Figure S1A; Table S1; 3,815 for young replicate 1; 3,569 for young replicate 2; 3,834 for aged replicate 1; 4,262 for aged replicate 2). Peak calling by MACS2 identified 478,861 chromatin accessibility peaks merged from all four samples distributed across distal (35%), intronic (53%), promoter (7%), and exonic (6%) regions (Figure S1B; Table S1). Clustering was performed using two modalities (RNA and ATAC) in an unbiased manner and resulted in 32 entities that recapitulated major cell types including excitatory neurons (ExN) in DG, CA1, 2, 3, and subiculum (SUB), and interneurons (ITN) including somatostatin- or parvalbumin-expressing interneurons (SST/Pvalb), Vip-expressing interneurons (Vip), Lamp5-expressing interneurons (Lamp5), astrocytes (ASTRO), neural stem cells (NSC), immature neurons (ImN), oligodendrocytes (OLIGO), oligodendrocyte precursor cells (OPC), microglia (MG), and endothelial cells (ENDO) (Figure 1B). The clusters were annotated by the composition of RNA expression and chromatin accessibility score (“gene score”), a predictor of gene expression based on accessible peaks surrounding the gene, for key cell type markers (Figure 1C–D; Figure S1C; Table S1). We validated our annotation for neuronal subtypes, clusters of excitatory neurons named DG (“*Prox1*”), CA1 (“*Mpped1*”), CA2 (“*Cacng5*”), CA3 (“*Cdh24*”), and SUB (“*Tshz2*”) by visualizing RNA expression and chromatin accessibility of their representative markers (Figure 1D–E).

**Figure 1.**
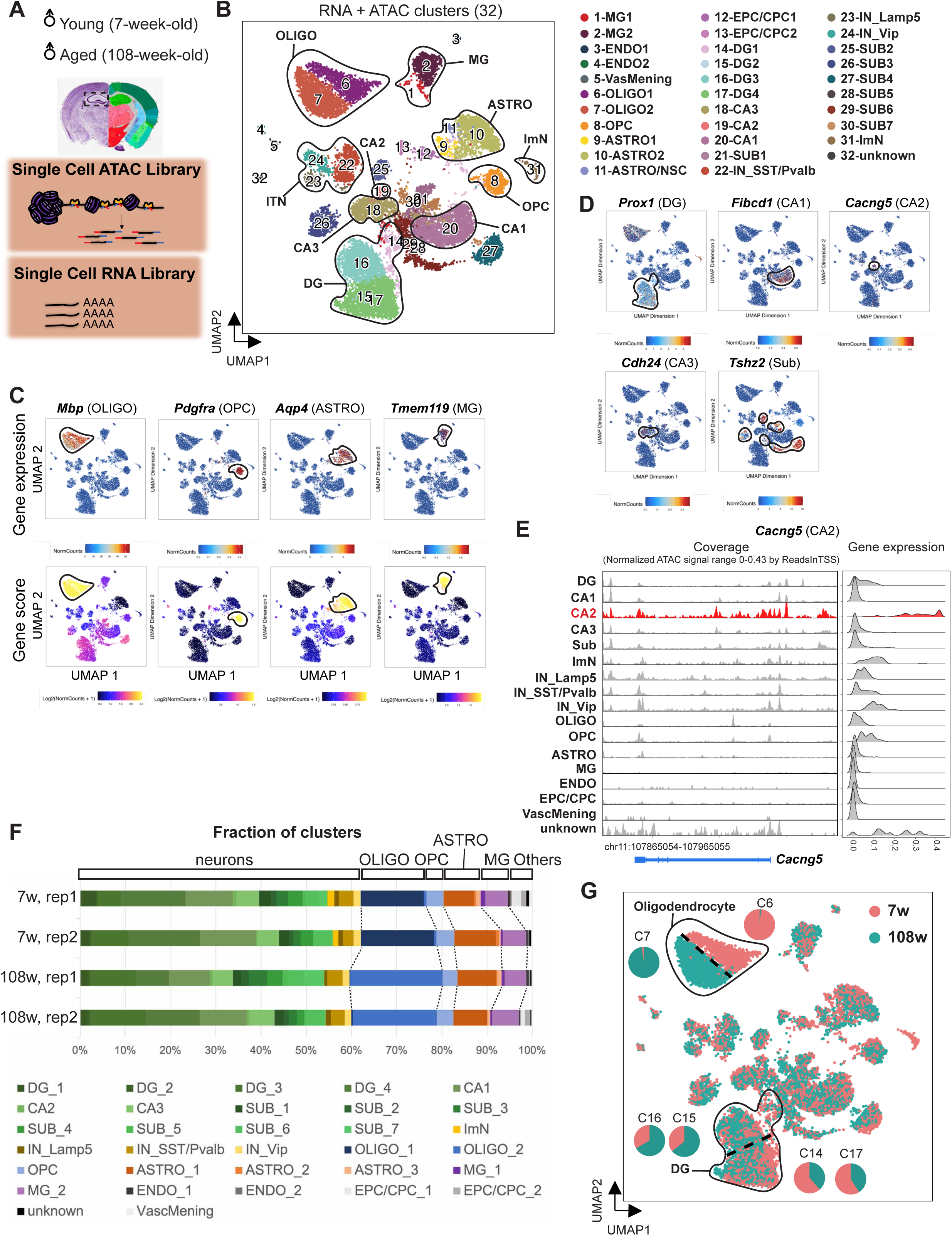
Single-nuclei profiling of transcriptome and chromatin accessibility in the aging mouse hippocampus. **A.** Scheme of experimental procedure for generating single-nuclei ATAC-seq and single-nuclei RNA-seq libraries using young (7-week-old) and aged (108-week-old) mice. Nissl and anatomical annotations were taken from the Allen Reference Atlas – Mouse Brain, atlas.brain-map.org. **B.** Visualization of clusters on UMAP generated by combining two modalities (RNA and ATAC). Cells are colored by clusters. **C–D**. Expression patterns of glial cell markers (C) and neuronal markers (D) are shown on UMAP. Clusters that strongly expressed these markers on the basis of heatmap intensity were encircled to highlight corresponding cell types. C. *Mbp*, *Pdgfra*, *Aqp4*, and *Tmem119* were used as representative markers for oligodendrocytes, oligodendrocyte precursors, astrocytes, and microglia, respectively. Gene expression (RNA assay, above) and gene score (ATAC assay, below) for these representative genes are shown. D. Gene expression (RNA) for neuronal markers is shown on UMAP. *Prox1*, *Fibcd1*, *Cacng5*, *Cdh24*, and *Tshz2* were used to represent DG, CA1, CA2, CA3, and SUB neurons, respectively. **E.** Genome browser track (left) and gene expression (right) for *Cacng5*, a representative marker of CA2 neurons. **F.** Percentage of clusters in B in two replicates from young and aged samples are shown. **G.** Visualization of cells on UMAP, colored by the age of sample collection. Age-dependent separation of OLIGO and DG is highlighted. Among OLIGO, 98.2% of cluster 6 consists of aged cells, while 97.7% of cluster 7 consists of young cells. Among DG, 66.7% and 64.1% of clusters 15 and 16 represent aged cells, while 61.4% and 59% of clusters 14 and 17 represent young cells.

Notably, similar to previous single-cell multiome studies on mouse skin, on human developing cortex, and on human peripheral blood mononuclear cells (Hao *et al*, 2021; Zhu *et al*, 2023; Ma *et al*, 2020), the integration of RNA and ATAC data provided a greater clustering resolution in cell type distinctions compared to a single modality (Figure S1E–F). For example, although the CA2 cluster with smaller cell numbers in the mouse brain was not distinguished on UMAP with the single RNA or ATAC modality, the integrative clustering revealed a clear separation between CA2 and CA3 neurons. Only RNA alone distinguished interneuron subtypes, whereas ATAC alone had ambiguous separation among excitatory neurons. ATAC clusters 11, 14, and 17 expressed DG, CA1, CA3, and SUB markers and did not correspond with a specific RNA cluster (Figure S1F). This approach was especially pivotal in delineating neuronal subtypes, and the joint profiling of chromatin accessibility and gene expression provided unique insights into excitatory neuron diversity (Figure 1B).

We confirmed the absence of over-clustering by assessing the separation of clusters according to the purity of the neighborhood for each cell as reflected by weighted high purity proportions (median > 0.9) (Methods; Figure S1D). Seurat’s label transfer method based on a prediction score using a spatial transcriptomics reference dataset of the hippocampus validated our cluster annotation for hippocampal neurons (Figure S1G; Table S1; Ortiz et al., 2020; Stuart et al., 2019). This prediction provided reliability of the identified neuronal subtypes, reaffirming the utility of integrating RNA and ATAC profiles for comprehensive cell type characterization, especially for neuron subtypes.

Our comprehensive profiling also illuminated significant age-dependent variations. Neuron populations dominated both young and aged samples by constituting over 60% of samples due to a nuclear isolation method allowing the capture of viable neurons (Figure 1F; Lake et al., 2016). We noticed that OLIGO was increased overall in frequency with aging (31% in young and 40% in aged) as also observed in other datasets (Allen *et al*, 2023), and significantly separated in an aged-dependent manner on UMAP (Figure 1F–G); clusters 6 and 7 were young- and aged-enriched, respectively. Similarly, aged and young DG neurons also had a tendency to group together (Figure 1G; Table S1); clusters 14 and 17 were young-enriched and 15 and 16 were aged- enriched.

Taken together, we generated single-nuclei transcriptome and open chromatin profiles of young and aged hippocampus from the same nuclei where the combination of these two assays resulted in a greater resolution of clustering and reflected the age-dependent separation of subtypes.

### Cell type–specific dynamics of transcriptome and epigenome during hippocampal aging

The aging-dependent segregation of OLIGO and DG clusters on UMAP implied that these cells might undergo more pronounced changes in gene expression and chromatin state with aging compared to other cell types. To assess age-related alterations specific to each cell type, we analyzed DEGs and DARs by comparing young and aged cells within each cell type (Methods).

We examined the aging-dependent transcriptomic changes in different neuron subtypes and in glial cells, including OLIGO (344 DEGs; adjusted *p*-value < 0.05) and ASTRO (85 DEGs; adjusted *p*-value < 0.05; Figure 2A; Table S2). Among excitatory neurons, DG neurons underwent the most pronounced transcriptomic changes with aging, followed by CA1, SUB, then CA3 neurons (DG: 169; CA1: 46; CA2: 0; CA3: 32; SUB: 43 DEGs; adjusted *p*-value < 0.05; Figure 2A; Table S2). At the chromatin level, neuron subtypes exhibited significant aging-associated changes, with a higher proportion of DARs compared to the total peaks detected than the proportion of DEGs to the total genes (Figure 2B–C). For example, the fraction of DARs in DG, CA1, and SST/Pvalb represented 0.92%, 0.48%, and 0.13% of total peaks, while the fraction of DEGs were 0.52%, 0.14%, and 0.02% among total genes, respectively. By contrast, glial cells showed a more balanced number of changes in their RNA and open chromatin profiles, indicating that both their transcriptome and chromatin accessibility were affected to a similar degree by aging (Figure 2C). Both DEG and DAR analyses did not detect major changes in ENDO and MG, which might reflect their insensitivity to aging-dependent alterations in their epigenome or transcriptome or technical issues as discussed later (Figure 2A–B). Notably, DG neurons and OLIGO exhibited the most drastic changes at both RNA and chromatin levels. Moreover, DEG analysis detected more aging-associated DEGs in non-neuronal cells, including OLIGO and ASTRO, than most neuronal subtypes, supporting previous observations (Allen *et al*, 2023).

**Figure 2.**
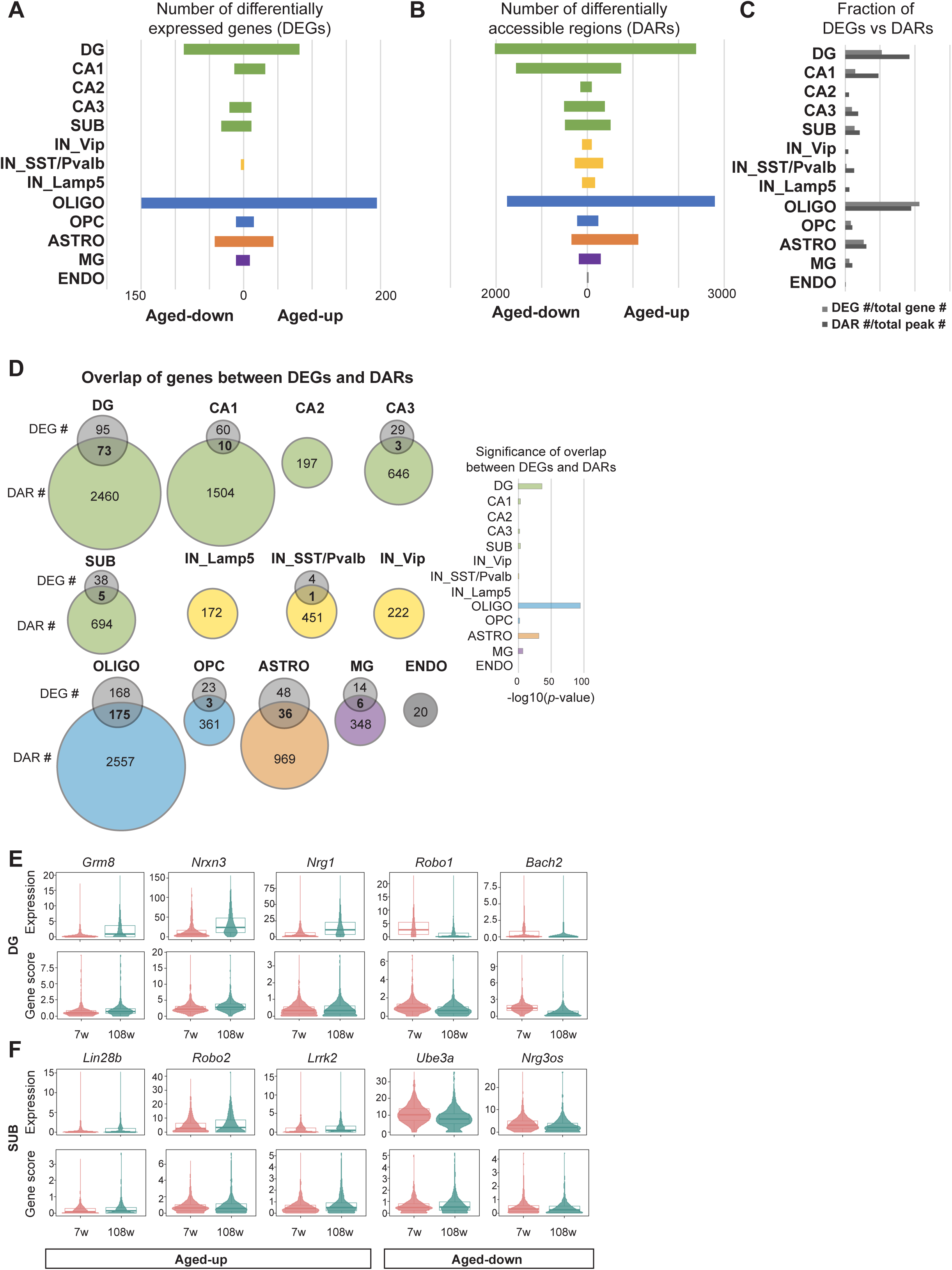
Cell type–specific transcriptome and epigenome dynamics across hippocampal aging. **A–B**. Number of DEGs (A) and DARs (B) by each major cell type. The graph is centered at 0 counts and divided into right and left directions to represent aged-upregulated and aged- downregulated genes or peaks, respectively. Color code of bars is determined by each major cell type. **C.** Bar plot showing the fraction of DEGs to total genes (light gray) and the fraction of DARs to total peaks (dark gray). **D.** Venn diagrams showing the number of genes obtained in DEG analysis (RNA) and DAR analysis (ATAC). Overlap of genes between DEGs and DARs are highlighted in bold. Significance of overlap was quantified by Fisher’s exact test and -log10(*p*-value) is plotted. *P*-values calculated for each cell type are as follows: DG, 2.4e-36; CA1, 1.5e-03; CA2, 1; CA3, 0.026; SUB, 2.3e-03; IN_Lamp5, 1: IN_SstPvalb, 0.068; IN_Vip, 1; Oligo, 6.9e-95; OPC, 3e-03; ASTRO, 5.1e-32; MG, 5.6e-08; ENDO, 1. **E–F**. Violin plots for gene expression (RNA) and gene score (ATAC) for DG-specific (E) or SUB- specific (F) representative overlapping aging genes in D.

**Figure 3.**
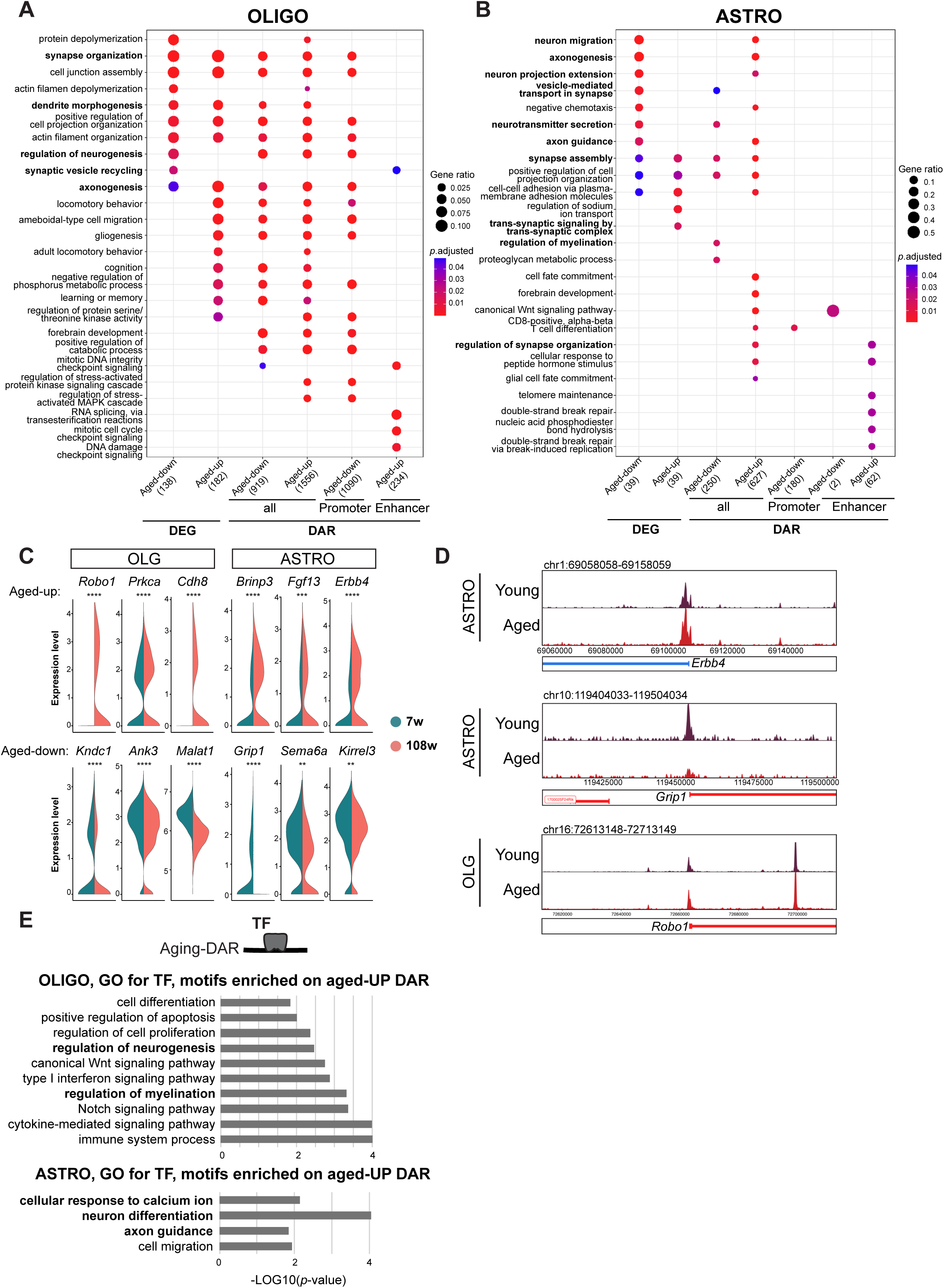
Dysregulation of neuronal genes in glial cells during hippocampal aging. **A–B**. Gene ontology enrichment analysis of DEGs, genes located near DARs, including all DAR peaks, peaks on promoters, and enhancers, separately. A, OLIGO. B, ASTRO. **C**. Violin plots for gene expression for synaptic function-related genes in OLIGO and ASTRO. Adjusted *p*-values obtained with Seurat’s *FindMarkers* function with the MAST algorithm using a hurdle model as a statistical test are shown (ns: non-significant; *: *p* < 0.05; **: *p* < 0.01; ***: *p* < 0.001; ****: *p* < 0.0001; Table S2). **D**. Genome browser tracks of chromatin accessibility near *Ebb4*, *Grip1*, and *Robo1*. Genes in the sense and antisense directions are shown in red and blue, respectively. The tracks are shown for young and aged cells separately. **E**. Bar plot of the enriched GO biological process terms for TF motifs enriched in aged-upregulated DARs of ASTRO and OLIGO.

The comparison of the fraction of annotated DARs altered during aging (aging DARs) revealed a notable enrichment of aging DARs in promoter regions in aged OLIGO and OPC compared to their young counterparts (Figure S2A; Table S2). In excitatory neurons (DG, CA1– 3, SUB), there was a global decline in distal intergenic peaks with aging and an increase in peaks on intronic and promoter regions in those cells, suggesting a shift in the regulatory landscape of these cells. Neuron subtypes, including DG, CA1, CA2, CA3, and SUB, displayed a higher number of nearest genes of aging DARs compared to the number of detected aging DEGs (Figure 2D). This result indicated that neurons underwent more profound changes at the chromatin level compared to the transcriptome level. Consistent with this, the nearest genes of DARs in CA1, CA2, CA3, and SUB neurons showed no or fewer overlap with DEG, while glial cells, such as OLIGO and ASTRO, had significant overlap (Figure 2D, Figure 4). This discrepancy in neurons suggests alterations in chromatin regulation in neurons that do not directly translate to gene expression changes, suggesting a chromatin state primed for aging features.

**Figure 4.**
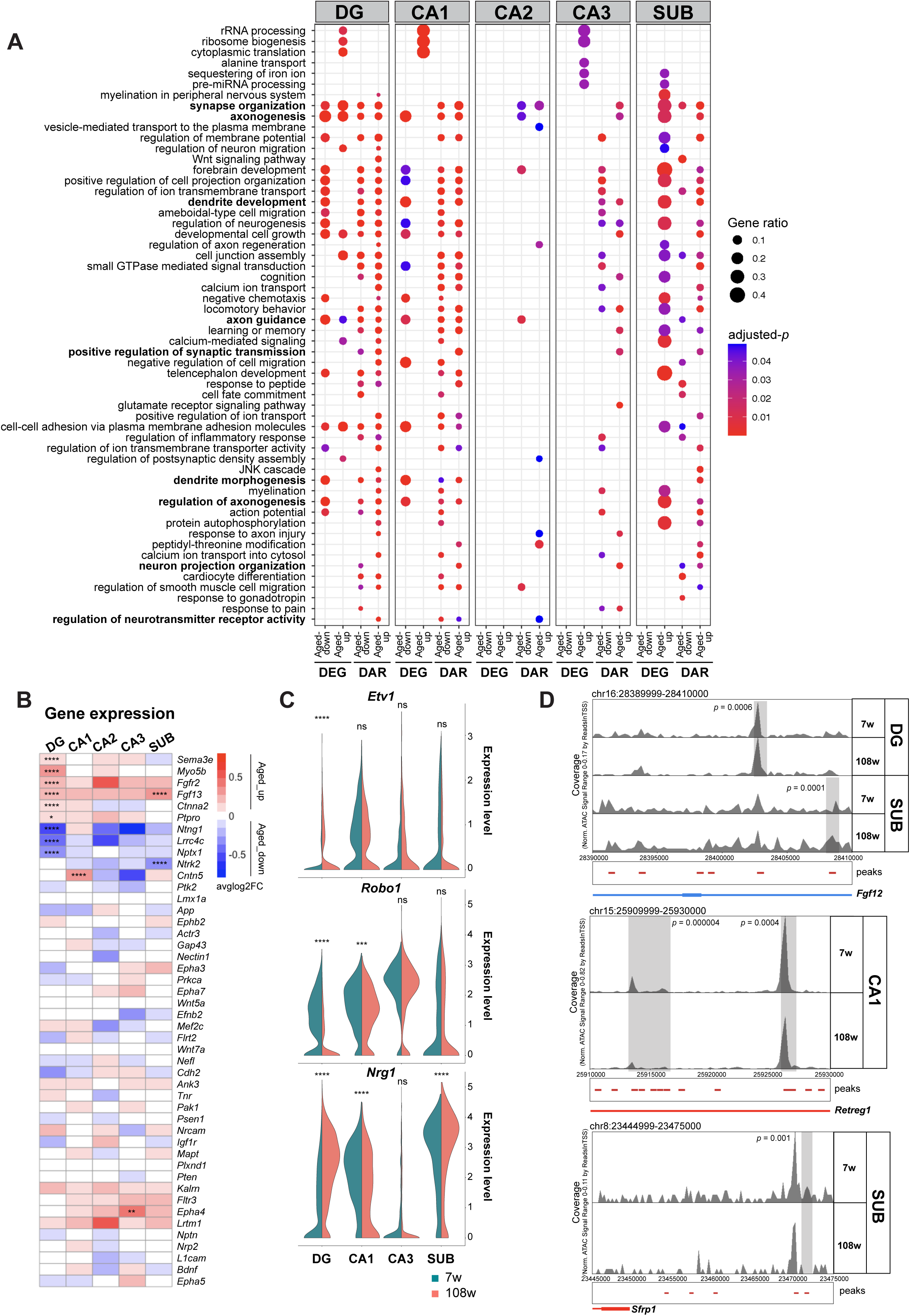
Chromatin accessibility–level dysregulations recapitulated aging features in neurons. **A.** Gene ontology enrichment analysis for biological processes in neuronal subtypes based on significant DEGs and nearest genes of DARs. Gene ratio is indicated by the size of circles and the color intensity indicates significance by adjusted *p*-value. **B.** Heatmap of averaged log-fold change of expression genes representative for axonogenesis- and synaptic organization–related terms in A, in young and aged cells of neuronal subtypes. Adjusted *p*-values obtained with Seurat’s *FindMarkers* function with the MAST algorithm using a hurdle model as a statistical test are shown (ns: non-significant; *: *p* < 0.05; **: *p* < 0.01; ***: *p* < 0.001; ****: *p* < 0.0001; Table S2). **C.** Violin plot of average gene expression for representative synaptic genes, *Robo1*, *Etv1*, and *Nrg1* in neuronal subtypes of young and aged mice. Adjusted *p*-values obtained with Seurat’s *FindMarkers* function with the MAST algorithm using a hurdle model as a statistical test are shown (ns: non-significant; *: *p* < 0.05; **: *p* < 0.01; ***: *p* < 0.001; ****: *p* < 0.0001; Table S2). **D.** Genome browser tracks of chromatin accessibility near *Fgf12*, *Retreg1*, and *Sfrp1*. Genes in the sense and antisense directions are shown in red and blue, respectively. The tracks are shown for young and aged cells separately. Black arrowheads indicate DARs between young- and aged- derived cells. Significant peaks are highlighted in gray. *P*-values obtained in edgeR’s DAR analysis are shown (Table S2; Methods).

To explore the aging-dependent gene regulatory mechanisms in different cell types, we have examined cell type–specific genes altered at both expression and chromatin levels. Notably, SUB neurons, a critical output structure of the hippocampus, have been relatively understudied in the context of aging. Our analysis identified specific dysregulation of synaptic genes within DG neurons, such as *Grm8*, *Nrxn3*, *Nrg1*, and *Robo1,* and immune-responsive transcription regulators, such as *Bach2*, suggesting the involvement of both synaptic and inflammatory components in aging (Figure 2E, Figure 5). In SUB neurons, *Lin28b*, *Robo2*, *Lrrk2*, *Ube3a*, and *Nrg3os* were among the dysregulated genes (Figure 2F). These genes play significant roles in various neuron functions, such as synaptic formation and axon guidance (Stolla *et al*, 2019; Blockus *et al*, 2021; Kuhlmann & Milnerwood, 2020; Furusawa *et al*, 2023; Müller *et al*, 2018). Their dysregulation sheds light on potential molecular mechanisms that might explain the vulnerability of SUB neurons, such as age-related decline in episodic memory (Chi et al., 2022). These findings highlight the specific molecular mechanisms potentially underlying aging-related vulnerability of SUB and DG neurons.

**Figure 5.**
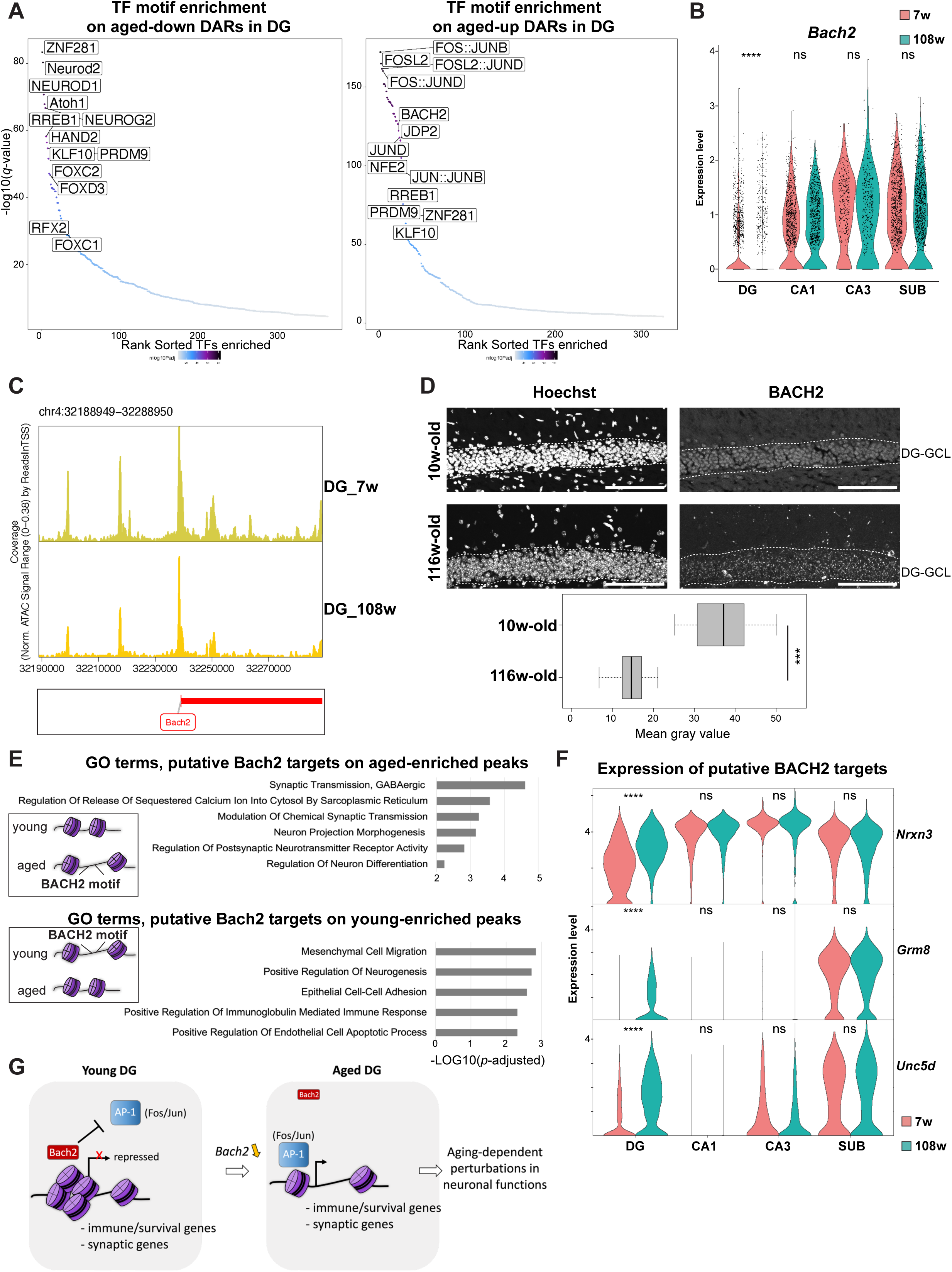
BACH2 activity reduced in aged DG neurons. **A.** TF motif enrichment analysis on aged-down DARs (left) and aged-up DARs (right) in DG neurons. The enriched TF motifs are distributed by their sorted rank on the *x*-axis enrichment and -log10(*q*-value) on the *y*-axis for significance. **B.** Violin plot of normalized gene expression for *Bach2* in each cell of neuronal subtypes split by their age. **C.** Genome browser tracks visualization of *Bach2* loci in DG neurons. The tracks are shown for young and aged DG cells separately. **D.** Immunostaining for anti-BACH2 in young (10w-old), and aged (116w-old) hippocampus. Images were cropped to DG area. Nuclei were counterstained with Hoechst 33342. Each box represents the distribution of mean gray value of the individual mice in each group (n = 2 mice for young; n = 3 mice for aged). Scale bars, 100 μm. ns: non-significant; *: *p* < 0.05; **: *p* < 0.01; ***: *p* < 0.001; ****: *p* < 0.0001. DG-GCL: dentate gyrus-granular cell layer. **E.** Gene ontology enrichment analysis for biological processes in DG neurons based on nearest genes of young and aged DARs with a BACH2 motif. The *x*-axis indicates significance by −log10(adjusted *p*-value). **F.** Violin plot of normalized gene expression for representative putative BACH2 targets, *Nrxn3*, *Grm8*, and *Unc5d*. Adjusted *p*-values obtained with Seurat’s *FindMarkers* function with the MAST algorithm using a hurdle model as a statistical test are shown (ns: non-significant; *: *p* < 0.05; **: *p* < 0.01; ***: *p* < 0.001; ****: *p* < 0.0001; Table S2). **G.** Model scheme for potential function of BACH2 in the aging process of DG neurons. Downregulation of BACH2 allows the aberrant activation of AP-1 target genes involved in immune response, survival, or synaptic function. This transcriptomic reprogramming might contribute to aging-associated functional decline in neurons.

To further examine the implication of AD risk genes among neuron subtypes with aging, we computed a module score using a set of genes altered during human AD progression based on a previous microarray analysis on the CA1 and CA3 regions (Table S2; Methods; Miller et al., 2013). Our analysis revealed that only DG and CA1 neurons exhibited significant changes with aging in the expression of genes downregulated or upregulated in AD, while other neuron subtypes did not show significant alterations (Figure S2). Similarly, at the chromatin accessibility level, aging DARs showed significant alterations in DG, CA1, and CA2 (Figure S2B). This suggests that both transcriptomic and chromatin accessibility changes in these AD risk genes are prominent in DG, CA1, and CA2 neurons.

Taken together, these results highlight the differential impact of aging across cell types and suggest a preparatory chromatin state for genes associated with neuronal dysfunction during aging. This state may contribute to the increased susceptibility of neurons to age-related stressors.

### Dysregulation of neuronal genes in glial cells during hippocampal aging

To elucidate the biological implications of the observed aging-related changes, we conducted gene ontology (GO) analyses for aging-associated DEGs and nearby genes of aging DARs, including promoters and enhancers (Methods). Several neuronal genes in glial cells, including OLIGO and ASTRO, were affected by the aging at both levels of gene expression and chromatin accessibility (Figure 3A–B; S3A; Table S3). For example, we observed that DARs related to vesicle-mediated transport in synapse and neurotransmitter secretion displayed decreased accessibility in ASTRO, showing a correlation with alterations in the transcriptome (Figure 3B). Specifically, the expression and chromatin accessibility of synaptic plasticity–related genes (e.g., *Cdh8*, *Prkca*, *Robo1, Kndc1, Ank3,* and *Malat1* in OLIGO; *Brinp3*, *Fgf13*, *Erbb4*, *Kirrel3*, *Sema6a*, and *Grip1* in ASTRO) were significantly altered with aging (Figure 3C–D). This suggests a dysregulated perisynaptic function involved in neuron–glia communication. A previous study performed oligodendrocyte-specific deletion of *Ank3* and revealed the function of glial AnkG, encoded by *Ank3*, on maintaining proper axoglial interactions during aging and contributing to behaviors of mice including motor learning, sociability, and depression (Ding *et al*, 2024). These findings also align with the potentially pivotal role of astrocytes in modulating the synaptic environment during aging, thereby influencing neuronal function and plasticity during aging (Popov *et al*, 2021).

Furthermore, enhancers associated with DNA integrity and mRNA splicing-related genes exhibited aging-related changes in OLIGO, while enhancers associated with telomere organization and double-strand break repair–related genes were affected in ASTRO (Figure 3A–B). These results suggest that global aging features, including mRNA splicing, telomere shortening, and altered DNA integrity, might be regulated at the enhancer level in glial cells.

To characterize the regulatory implications of these DARs, we performed a motif enrichment analysis and a GO analysis using all human transcription factors as a background (Figure S3B, Table S3). Neuronal function–associated terms were listed (Figure 3E). Specifically, regulation of neurogenesis was enriched in OLIGO, while neuron differentiation, cellular response to calcium ion and axon guidance were enriched in ASTRO. These findings suggest a significant reconfiguration of gene regulatory programs in glia, highlighting their potential role in contributing to the functional decline observed in aging neurons.

### Chromatin accessibility–level dysregulations recapitulated aging features in neurons

We next examined aging-associated dysregulations at chromatin accessibility and gene expression levels in neurons. GO term analysis unveiled common features shared by neurons and glial cells, notably in synapse organization and axonogenesis (Figure 3, 4A; Table S4). Among neurons, DAR analysis using all aged-up and aged-down peaks revealed terms consistently detected across major cell types, including synapse organization, axonogenesis, learning, and cognition, which were not necessarily enriched in DEG analysis in all neuronal types (Figure 4A–B). Some of the neuronal genes whose expression was commonly altered with aging included *Robo1* and *Nrg1*, or specifically altered in DG (e.g. *Etv1*), which are involved in axon guidance and synaptic plasticity (Figure 4C). *Robo1* was previously shown to exhibit reduced chromatin accessibility in their putative enhancers and promoters with hippocampal aging (Figure 2E; Zhang et al., 2022).

Given that DEG analysis did not detect terms associated with synaptic plasticity, particularly in CA1, CA2, and CA3 neurons (Figure 4A-B; Table S4), these findings indicate that aging-related transcriptomic changes only partially recapitulate aging features observed *in vivo* including axonogenesis and dendrite morphogenesis, while chromatin accessibility changes could provide a more comprehensive picture on other neuronal terms, such as synapse organization, synaptic transmission, learning or memory, and neurotransmitter receptor activity. These cells appear to undergo profound chromatin remodeling with aging, which is not reflected in gene expression in the steady state, which we analyzed in this study. This preparatory chromatin state may predispose these neurons to age-related functional decline and increased susceptibility to age-associated diseases. This phenomenon was also observed in immune genes, exemplified by a primed chromatin state in oligodendrocytes of control samples in the context of multiple sclerosis (Meijer *et al*, 2022). We further confirmed the absence of synaptic plasticity–related terms in GO analysis using aging DEGs in excitatory neurons derived from publicly available single-cell, not single-nucleus, RNA-seq datasets. This approach helped avoid the potential issue of a lack of neuronal mRNA species due to single nuclei isolation for sequencing rather than whole cells (Table S4; Methods; Ogrodnik et al., 2021). We found that, rather than synaptic function–related genes, inflammation and immune response genes were dominant among aging DEGs (Figure S4B). Taken together, these data suggest a preparatory chromatin state for neuronal function–related loci during normal aging, indicating that hippocampal neurons may be predisposed to age-related brain dysfunctions. This preparatory state could make these neurons more susceptible to aging and aging-associated diseases when exposed to additional stressors.

Additionally, GO analysis for chromatin accessibility showed distinct alterations in specific cell types, revealing unique vulnerabilities during aging (Figure 4A). Notably, SUB and DG neurons exhibited increased accessibility in gene loci associated with JNK cascade (e.g., *Fgf12*), involved in axonal response and synapse growth, highlighting their distinctive susceptibility to aging (Figure 4D; Etter et al., 2005; Raivich et al., 2004). CA1 neurons displayed an altered transcriptome related to rRNA metabolism, dendrite development, and axon guidance. On the other hand, a putative enhancer and the promoter of *Retreg1*, known for its neuroprotective role (Mo *et al*, 2020), were less accessible in aged CA1 neurons, but the expression was not altered (Figure 4D, S4C). SUB neurons exhibited alterations in potential regulator loci outside gene associated with Wnt signaling (e.g., *Sfrp1*), indicating their specific cellular responses to aging stressors (Figure 4D).

Together, these results highlight cellular processes affected by aging, emphasizing both shared and cell type–specific vulnerabilities among neurons. These findings underscore the importance of chromatin-level changes, which are potentially susceptible to driving the functional decline observed in aging neurons.

### Bach2 activity reduced in aged DG neurons

We then focused on DG neurons as they exhibited the most significant transcriptomic and chromatin level changes with aging among all neuron subtypes, including neuronal function– related genes (Figure 4A). To uncover cis-regulatory mechanisms underlying DG aging, we performed motif-enrichment analysis on aging DARs, revealing a pronounced enrichment for motifs associated with neuron differentiation, axon development, axon guidance, and cellular response to calcium ions (Figure S5A–B; Table S5). Whereas aged-accessible DARs in DG neurons were enriched in motifs of TFs consisting of the activator protein-1 (AP-1) complex, such as JUN family proteins, FOS, and FOSL2, and the BTB and CNC Homology (BACH) family, young accessible DARs were enriched in NEUROD1/2 and NEROG2 that are involved in neurogenesis (Figure 5A). The AP-1 complex, composed of cJUN, JUNB, JUND, cFOS, FOSB, FOSL1, FOSL2, BATF, and BATF3, can bind to Maf recognition elements to activate the expression of the target genes (Eferl & Wagner, 2003; Shaulian & Karin, 2002). By contrast, BACH1 and BACH2 transcription factors recognize a similar motif for repression in a competitive manner with the AP-1 complex (Jang *et al*, 2017; Itoh-Nakadai *et al*, 2014; Tsukumo *et al*, 2013). The AP-1 complex plays a critical role in regulating gene expression in response to a variety of physiological and pathological stimuli, such as neuronal activation, and is involved in synaptic plasticity, memory formation, and response to stress (Alberini, 2009; Sanyal *et al*, 2002). In these studies, while the function of the AP-1 complex is well-documented in neurons, the role of BACH2 is not well understood; its role in DNA damage, apoptosis, immune response, and proliferation has been documented in other contexts such as immune cells (Uittenboogaard *et al*, 2013; Liu & Liu, 2022).

Notably, we observed a decreased expression of *Bach2* specifically in DG neurons, which exhibited the most notable changes in transcriptome among all TFs enriched on aging-up DARs (Figure 5B; S5D). We hypothesized that the reduction of BACH2, a transcription factor involved in cellular stress responses, contributes to the decline in neuronal function and neuron death in DG neurons. The accessibility of the BACH2 promoter, gene body, and putative enhancers also diminished in aged DG neurons (Figure 5C). We confirmed the decline of BACH2 expression in the aging hippocampus *in vivo* by immunohistochemistry (Figure 5D). The GO analysis identified an enrichment of genes involved in synaptic transmission (e.g., *Nrxn3* and *Grm8*), and apoptosis (e.g., *Unc5d*) in aged-accessible DARs in DG neurons with BACH2 motifs (Figure 5E–F; Table S5). The expression of these putative target genes was significantly upregulated in DG neurons (Figure 5F; Figure S5C). While the overall expression of the nearest genes of the BACH2 motif– containing aged-accessible DARs averaged in each neuron subtype significantly increased in aged DG neurons, we observed the opposite tendency for the nearest genes of young-accessible DARs (Figure S5C). This suggests a dual and selective role for BACH2 in activating and repressing the downstream target genes or a complementary effect of other factors, such as the AP-1 complex.

These findings suggest a role for *Bach2* in the epigenetic and transcriptional landscape of DG neurons during aging. The diminished *Bach2* expression and its motif activity on aging DARs highlight its regulatory role over genes critical for synaptic organization, apoptosis, and immune response. The upregulation of putative target genes in aged DG neurons signifies a potential mechanism through which BACH2 influences aging-associated neuronal functions. Taken altogether, these results raise the possibility that the decreased *Bach2* expression in aged DG neurons induces de-repression of its target genes and contributes to aging-associated functional decline in DG neurons by controlling the expression of synaptic and apoptotic genes (Figure 5G).

## DISCUSSION

In this study, we systematically characterized cell type–specific gene regulatory networks in the aging mouse hippocampus using single-nucleus multiome sequencing. Our findings revealed significant age-dependent changes in both transcriptome and chromatin accessibility across various cell types, highlighting differential vulnerabilities particularly among DG, CA1, and SUB neurons, as well as glial cells. Notably, we observed substantial variations in the composition of oligodendrocytes and DG neuron clusters with age, and pronounced transcriptomic and chromatin accessibility alterations were seen in these cell types.

We observed greater chromatin-level changes in both glial cells and excitatory neurons (DG, CA1, CA2, CA3, and SUB) than transcriptome-level changes, underscoring the critical role of chromatin remodeling in aging-associated functional decline. The absence of immediate transcriptomic alterations in synaptic plasticity–related genes suggests the existence of a preparatory chromatin state for genes associated with neuronal dysfunction during aging. While this chromatin priming may not translate into immediate changes in gene expression during normal cognitive aging, it could become significant when neurons are subjected to vulnerable stressors, contributing to memory impairment or the progression of neurodegenerative diseases. These findings imply that changes in the chromatin state, rather than immediate changes in gene expression, may underpin susceptibility to aging and neurodegenerative diseases. This preparatory state may be partly explained by the loss of heterochromatin domains with aging, as previously reported in excitatory neurons (Zhang *et al*, 2022). Future research should explore epigenetic marks that could contribute to altered chromatin accessibility during aging.

Previous bulk transcriptomic and microarray studies have revealed a global decrease in neuronal genes, particularly those associated with synaptic function–associated genes, and an increase in immune genes in the aging brain (Ham & Lee, 2020; Loerch *et al*, 2008; Lu *et al*, 2004; Berchtold *et al*, 2008; Erraji-Benchekroun *et al*, 2005; Bae *et al*, 2018; Naumova *et al*, 2012; Berchtold *et al*, 2013). However, these studies did not differentiate age-associated gene expression changes based on neuronal or glial origin, as the profiling was not restricted to specific cell types. Recent single-cell studies, including the current one, indicated that not all excitatory neuron subtypes exhibit alterations in neuronal function–related genes in their transcriptome with aging (Allen *et al*, 2023; Morabito *et al*, 2021). While the neuronal subtype distinction was not specifically addressed in these studies, our study has provided neuron subtype–specific alterations in gene expression and chromatin accessibility in the aging hippocampus. Among neurons, only DG and SUB neurons, along with glial cells, undergo significant changes due to alterations in synaptic genes. This finding contrasts with the bulk studies, suggesting that the effects of aging on gene expression are more cell type–specific. Moreover, previous research on bulk studies reported alterations of synaptic plasticity–related genes at the mRNA splicing level rather than the expression level (Stilling *et al*, 2014; Tollervey *et al*, 2011). Future studies should investigate the cell type–specific changes in mRNA splicing in the aging context and explore the relationship between chromatin state and post-transcriptional regulations (e.g., mRNA splicing) for these genes in normal cognitive aging and neurodegenerative diseases. Our findings thus suggest that chromatin accessibility rather than gene expression recapitulates synaptic plasticity–related dysregulations in a cell type–specific manner during hippocampal aging.

Notably, our study identified AP-1 and BACH2 as potential critical factors in the aging-mediated functional decline of DG neurons, characterized by increased expression of *Jun* and decreased expression of *Bach2*, and heightened motif activity for both with aging (Figure 5A–B; S5D). Our findings align with recent studies that report an enhanced activity of the JUN motif in AD excitatory neurons, as well as increased activity of JUNB and cJUN motifs in DG neurons with aging (Zhang *et al*, 2022; Morabito *et al*, 2021). BACH2, known for its role in regulating immune response and cellular stress, has been shown to decrease in expression with age in various tissues including in healthy human lymphocytes and lung, liver, kidney, and spleen of aged mice. Conversely, AP-1 underwent activation with inflammation and its motif is enriched in aged neuronal lineages (Long *et al*, 2019; Karakaslar *et al*, 2023; Uittenboogaard *et al*, 2013; Zhang *et al*, 2022). In this study, we observed the downregulation of BACH2 and the enrichment of AP-1 and BACH2 motifs in aging DARs within DG neurons, which suggests BACH2’s involvement in regulating genes associated with neurogenesis, immune response, synaptic organization, and apoptosis. Notably, aging of pyramidal neurons is linked to increased excitability due to changes in membrane properties and synaptic function (Simkin *et al*, 2015; Vaillend *et al*, 2002). Additionally, aged memory-impaired animals display enhanced cFOS activity in response to neural stimulation (Haberman *et al*, 2017). BACH2 may regulate the activity of cFOS, a component of the AP-1 complex, by competing in transcription regulation. As BACH2 levels decline with age, this reduction may allow activation of immediate early gene, contributing to altered intrinsic neuronal excitability. Our findings provide new insights into the potential molecular mechanisms underlying aging-associated cognitive decline and highlight BACH2 as a promising therapeutic target for improving hippocampal function in aging and neurodegenerative diseases.

Finally, our dataset exhibited limitations in detecting immune-related genes due to the profiling of nuclei instead of cells in most cell types, particularly in microglia. This result aligns with previous findings of the differential capacity of mRNA capture between nuclear and cellular compartments of microglia in single-cell studies. Specifically, single-nuclei transcriptome sequencing has been shown to lack microglia activation genes, including many immune-related genes, as compared to single-cell sequencing (Thrupp *et al*, 2020). This underscores the need for technical considerations when interpreting immune-related gene expression in nuclear versus cellular transcriptomics.

In summary, this study provides a comprehensive overview of the cell type–specific dynamics of transcriptome and epigenome during hippocampal aging. Our findings highlight the unique vulnerabilities of distinct cell types and emphasize the importance of chromatin-level changes in understanding the aging process. Identifying critical factors, such as BACH2, and elucidating their roles in age-related changes, paves the way for future research to unravel the mechanisms of age-related cognitive decline and develop targeted interventions for preserving cognitive function in the aged populations.

## MATERIALS AND METHODS

### Ethics statement

All animals were housed and studied in compliance with protocols approved by the Animal Care and Use Committee of The University of Tokyo. The approval numbers are P25-8 and P30-4 from the Graduate School of Pharmaceutical Sciences, and 0421 and A2022IQB001-06 from the Institute for Quantitative Biosciences. All procedures were followed in accordance with the University of Tokyo guidelines for the care and use of laboratory animals and ARRIVE guidelines.

### Mouse maintenance and brain tissue samples

Wild-type C57BL/6N mice were purchased from CLEA Japan. Some of the aged mice were provided by the Foundation for Biomedical Research and Innovation at Kobe through the National BioResource Project of the Ministry of Education, Culture, Sports, Science and Technology (MEXT), Japan. All mice were maintained in a temperature- and humidity-controlled environment (23 ± 3°C and 50 ± 15%, respectively) under a 12-h light/dark cycle. Animals were housed in sterile cages (Innocage, Innovive) containing bedding chips (PALSOFT, Oriental Yeast), at a density of two to six mice per cage, and provided irradiated food (CE-2, CLEA Japan) and filtered water ad libitum. Animals used for sn-multiome-seq were euthanized at 7 weeks and 108 weeks of age, with two males used at each age.

For validation with immunohistochemistry, mice were euthanized at 10 weeks, and 116 weeks. Mice were anesthetized with isoflurane and perfused with ice-cold PBS followed by 4% paraformaldehyde (PFA) in PBS. Brains were postfixed in 4% PFA overnight at 4°C, rinsed in PBS and incubated at 4°C in 10%, 20%, and 30% sucrose until they sank to the bottom of the tube before being frozen in optimal cutting temperature (O.C.T.) compound (Sakura) and stored at −80°C.

### Library preparation for single nuclei multiome (RNA+ATAC) sequencing

The protocol for nuclear isolation from brain tissues was adapted from a previous study (Frey *et al*, 2023; Bundo *et al*, 2016). All procedures were carried out on ice. Frozen hippocampi of young and aged mice were sequentially homogenized using a syringe with 23G and 27G needles in 500 μL of 54% Percoll in homogenizing buffer (50 mM Tris-HCl pH 7.4, 25 mM KCl, 5 mM MgCl2, and 250 mM sucrose). The homogenate was then mixed with 10% NP-40 (final concentration, 0.1%), incubated on ice for 15 minutes, and mixed with 500 μL of homogenizing buffer. Then, Percoll gradient was prepared in the following order: 100 μL of 35% Percoll in homogenizing buffer at the bottom layer, 200 μL 31% Percoll in the middle, and 1 mL of homogenate (27% Percoll) on the top layer. The tube was centrifuged at 20,000 × *g* for 10 minutes at 4°C. After removing the debris from the top layer, nuclei were collected from the bottom layer and transferred to a new tube. The nuclear pellet was resuspended in PBS containing 0.02% BSA. The nuclei were counted and diluted to a concentration of 4,000 nuclei per microliter in PBS with 0.02% BSA.

The nuclei were then used for library preparation, targeting the recovery of 4,000 nuclei per sample using a chromium system (10x Genomics, Next GEM Single Cell Multiome ATAC + Gene expression Reagent Bundle, PN-1000283). The libraries were sequenced on the DNB-seq platform to obtain 100-base paired-end reads.

### Alignment, raw processing, and quality control of sc-multiome-seq data

Fastq alignment to the reference genome (mm10), barcode processing, single-cell counting, and read quantification and filtering were performed on each sn-multiome-seq library using the Cell Ranger-Arc pipeline version 2.0 (10x Genomics). Intronic reads for all samples were included to quantify pre-mRNA transcripts to account for unspliced nuclear transcripts. Cell Ranger-Arc filtered count matrices were used for downstream analysis.

### Processing, cell clustering, visualization, and cluster annotation

ArchR version 1.0.2 was used for processing the paired snRNA-seq gene expression data and snATAC-seq fragment data for all samples (Granja *et al*, 2021). ArchR framework was used for quality control filtering steps, single-cell clustering, peak calling, and differentially accessible region analysis. ATAC fragments were read using the *createArrowFiles* function. Gene expression values were added using the *addGeneExpressionMatrix* function. Nuclei were excluded from downstream analysis if the TSS enrichment score was < 6, fewer than 3,000 unique nuclear fragments, and missing matched RNA reads. Nuclei doublets were excluded with the *addDoubletScores* and *filterDoublets* functions of ArchR using TileMatrix of the snATAC-seq. The removal of doublets resulted in filtering of 4% and 3.7% of two respective replicates of 7- week-old samples, and that of 4% and 4.4% of two respective replicates of 108-week-old samples.

As a result, 15,480 nuclei were used for downstream analysis with a median TSS score of 13,441 and median fragments per nucleus of 27,484.5.

Dimension reduction was performed using each GeneExpressionMatrix derived from snRNA-seq and TileMatrix derived from snATAC-seq using the *addIterativeLSI* function of ArchR with default parameters and resolution set to 0.2 for clustering. We used the *addCombinedDims* function to combine two-modality (RNA and ATAC) dimension reductions into a single reduction. For each reduction, Uniform Manifold Approximation and Projection (UMAP) was performed with default parameters of the *addUMAP* function. Cell clusters were generated using Seurat-implemented in ArchR using the combined RNA and ATAC matrix with a resolution set to 0.8. Cell type annotation was performed on the basis of gene scores and gene expression. Gene score represents gene expression and is calculated on the basis of chromatin accessibility at the gene body, promoter, and distal regulatory regions. We identified marker genes for each cluster using the *getMarkerFeatures* function using GeneScoreMatrix and GeneExpressionMatrix (Table S1). We further confirmed cluster identities by following known markers for each major cell type: *Snap25*, *Syt1*, and *Celf4* for general neuron markers; *Slc17a7* and *Satb2* for excitatory neurons; *Prox1* for DG neurons; *Mpped1*, *Wfs1*, and *Dcn* for CA1 neurons; *Sulf2* and *Nrip3* for CA3 neurons; *Dlx1* and *Gad1* for interneurons; *Sst*, *Vip*, *Lamp5*, and *Sncg* for interneuron subtypes; *Cldn11*, *Plp1*, and *Mbp* for oligodendrocytes; *Olig1*, *Pdgfra*, and *Sox10* for oligodendrocyte precursors; *Slc1a3*, *Slc1a2*, *Gjb6*, *Glul*, *Aqp4*, *Fgfr3*, *Aldh1l1*, and *Gfap* for astrocytes; *Cx3cr1* and *Itgam* for microglia; *Col1a2* and *Slc6a13* for vascular meningeal cells; *Flt1*, *Pecam1*, and *Cldn5* for endothelial cells; and *Ttr* for ependymal or choroid plexus cells. Next, we merged clusters displaying similar markers for DEG and DAR analyses.

Seurat version 4 was used for differential gene expression analysis on a total of 14,908 nuclei carried over from ArchR object. Merged Seurat object containing ArchR-filtered cells was log-normalized and scaled with default parameters. Eighteen principal components and 0.8 of resolution were used for clustering and UMAP reduction using the *FindNeighbors* and *RunUMAP* functions of Seurat. We confirmed that the median of the percentage of mitochondrial transcript in all carried-over nuclei was 0.062, representing the healthy state of the samples.

### Peak Calling and annotation

Pseudo-bulk replicates were generated for major cell types using the *addGroupCoverages* function of ArchR. The reproducible peak sets for each major cell type at different ages were named with default parameters using the *addReproduciblePeakSet* function implementing MACS2 (version 2.2.5) to create a fixed-width peak size of 501 bp and iterative overlap peak merging on the basis of coverage data grouped by each major cell type split by age. A total of 478,861 peaks were generated and used for downstream analysis. Peaks were required to be present in at least two pseudo-bulk replicates. Using mm10 annotations from BSgenome.Mmusculus.UCSC.mm10 package version 1.4.3 in R, we annotated the peak set of each cell type from young and aged samples into the following categories: promoter, exonic, intronic, and distal (Table S1). For the annotation of aging DAR peaks, we have used the *annotatePeak* function of the ChIPSeeker package in R (Table S2; Yu et al., 2015). For the annotation of promoter and enhancer regions in aging DAR peak lists, *bedtools intersect* was used (Quinlan & Hall, 2010). For promoters, we have used the promoter list generated by the Cellranger-Arc pipeline. For enhancers, we have withdrawn the file of enhancer-gene interactions identified in the cortex from http://www.enhanceratlas.org/downloadv2.php.

### Differentially accessible region analysis and finding nearest genes

EdgeR was employed to generate differentially accessible regions in each of the major cell types between young and aged cells (Chen *et al*, 2024). Cell type–focused analysis rather than subtype (cluster)-focused one was chosen to increase the coverage of clusters. The LRT method was used to obtain reliability of two replicates. DARs were considered significantly different if they had a *p*-value < 0.001 in downstream analyses (Table S2).

Aging DARs were annotated using the *annotatePeaks* function of the ChIPSeeker package with default parameters (Yu *et al*, 2015). The column *geneId* of the output file was used to define nearest genes (Table S2).

### TF motif enrichment analysis

TF motif enrichment analysis on aging DARs was performed using Simple Enrichment Analysis (SEA) version 5.5.2 (Bailey & Grant, 2021). Bed files containing significant aging DARs obtained as described above were converted to fasta format using bedtools’ getfasta, to use as input on SEA. Motif annotations were obtained using JASPAR CORE (2022) vertebrates database with default parameters. The TF motifs were considered significantly enriched if the *E*-value (multiplication of *p*-value by the number of motifs in the input) was < 10. The enriched TF motifs are ranked according to their *E*-value. All motifs in the output files from SEA had a *q*-value < 0.01 (Table S3–S4).

### Annotation of putative BACH2-target genes

FIMO version 5.5.0 was used as a motif search tool to find aging DARs containing the MA1101.2.meme motif (Grant *et al*, 2011). A total of 2,383 sequences and 2,021 sequences identified in the DAR analysis for DG neurons were applied for aged-enriched and young-enriched peaks, respectively. We used bedtools to annotate the genomic regions with the motif. This annotation identified 546 and 245 genes for aged- and young-enriched peaks, respectively (Table S5). The resultant gene list was applied to the enrichR web (Chen *et al*, 2013). We queried GO Biological Process 2023 (Table S5). The top 5 terms (ranked by FDR) are displayed.

### Module score calculation for Alzheimer’s disease genes and BACH2-putative target genes

To calculate the average expression levels of Alzheimer’s disease genes and BACH2-putative target genes at a single-cell level, we applied the *addModuleScore* function of ArchR implementing the Seurat *addModuleScore* function on GeneExpressionMatrix for gene score or GeneExpressionMatrix for gene expression. We set the background number as 5 genes for signal normalization and bins to 25 for background calculation. This function generated a module score by calculating the average expression of each test gene group, subtracted by the aggregated expression of control gene sets. The Kruskal–Wallis test was performed for significance (ns: non- significant; *: *p* < 0.01; **: *p* < 0.001; ***: *p* < 0.0001).

### Differential gene expression analysis

ArchR cluster assignments were added to Seurat metadata. The merged Seurat object was normalized to the sequencing depth and scaled with default parameters. DEG analysis was performed on ArchR clusters by the *FindMarkers* function in Seurat using the two-hurdle model in DE testing implemented in the MAST algorithm to reduce false positives. A cell type–focused analysis rather than subtype (cluster)-focused one was chosen to increase the coverage of clusters. To regress out technical biases from different samples, the latent.vars = “batch” option was used to perform DE testing. Log2(fold-change) values of the average gene expression and adjusted *p*- values were generated between young- and aged-derived cells of each major cell type.

The list of DEGs for major cell types was generated including all detected genes (Table S2). For downstream analyses, such as Gene Ontology analysis, the genes were filtered for adjusted *p* < 0.05.

### Overlap of DEGs and nearest genes of DARs

Statistical significance of the overlap of genes between DEGs and nearest genes of DARs was computed by Fisher’s exact test using the GeneOverlap package in R (v1.38). The background for the gene overlap analysis was defined as the total number of genes detected in the merged Seurat object.

### Gene ontology term enrichment analysis

To identify the biological processes of each list of nearest genes of DARs, promoters and enhancers on DARs, and DEGs in indicated cell types, clusterProfiler version 4.2.2 in R was used as an enrichment tool (Table S3–S4; Wu et al., 2021; Yu et al., 2012). Enrichplot version 1.14.2 in R was used to visualize functional enrichment results. The top five terms with distinct gene sets are displayed (ranked by FDR).

For DEGs identified in the Ogrodnik dataset, the list of significant DEGs (adjusted *p* < 0.05) was applied to the enrichR web (Table S4; Chen et al., 2013).

For TFs, DAVID was used with the background defined as 1,639 human TFs (Lambert *et al*, 2018; Huang *et al*, 2009a, 2009b).

### Cluster purity analysis

Bluster package in R was used to assess the cluster separation. Cluster purity of ArchR clusters was computed with the *neighborPurity* function in the UMAP expression space. Median purity greater than 0.9 indicated that cells from the same cluster are mainly surrounded by the other cells in the same cluster, reflecting that clusters are well-separated.

### Processing of external single-cell RNA-seq data sources

To validate our DEG analysis on single-nuclei RNA-seq, we analyzed previously published single-cell RNA-seq studies of the aging brain. For the Ogrodnik dataset (Ogrodnik et al., 2021), two replicates of young and aged single-cell RNA-seq samples were withdrawn from the accession number GSE161340 of the NCBI Geo repository and merged in Seurat. Cells with more than 200 and less than 5,000 genes were removed. Twenty-five principal components were used for clustering and UMAP dimension reduction in the *FindNeighbors* and *RunUMAP* functions. The resolution was set to 0.8 in the *FindClusters* function, generating 23 clusters consisting of microglia, oligodendrocytes, endothelial cells, microglia, astrocytes, oligodendrocyte precursors, dentate gyrus neurons (DG), CA1 neurons, immature neurons, pericytes, choroid plexus cells, vascular cells, inhibitory neurons, and unknown cells. DG and CA1 clusters were further subsets for DEG analysis. The *FindMarkers* function was applied between young and aged cells of the major cell types for DE testing with default parameters. Significant DEGs with adjusted *p* < 0.05 were used for downstream analyses (72 genes).

For the Ortiz dataset (Ortiz et al., 2020), raw expression data and original metadata were withdrawn from accession number GSE147747 of the NCBI Geo repository. Hippocampal cells were subset using CA1slm, CA1so, CA1sp, CA1sr, CA2slm, CA2so, CA2sp, CA2sr, CA3slm, CA3slu, CA3so, CA3sp, CA3sr, DG-mo, DG-po, DG-sg, SUBd-m, SUBd-sp, SUBd-se, SUBv-sr, SUBv-m, SUBv-sp, SUBv-sr, PAR1, PAR2, PAR3, POST1, POST2, POST3, PRE1, PRE2, and PRE3 clusters. We performed sctransform-based normalization. We then performed the transfer labels method of Seurat by applying the *FindTransferAnchors* and *TransferData* functions using our dataset subset with neuronal clusters with sctransform-based normalization, which identified 587 anchors.

### Immunohistochemistry and confocal imaging

Serial 14 µm coronal cryosections were cut on a Cryostat (TissureTek) and stored at −80°C until use. The sections were washed in 0.1 Tween-20 in TBS, and blocked in 0.1 Triton X-100 in TBS. The sections were stained with anti-Bach2 (rat; Abcam, ab243148, 1:250 dilution), anti–NeuN- 488 (mouse; Millipore, MAB377X, 1:500 dilution) overnight at 4°C, followed by staining with donkey anti–rabbit Alexa Fluor 647 (Invitrogen, A32795, 1:1,000 dilution) and Hoechst (Invitrogen, H3570, 1:10,000 dilution), for 1 hour at room temperature. Primary antibodies were used in blocking buffer, and secondary antibodies and Hoechst were diluted in TBS with Tween-20. Slides were mounted with the mounting medium before imaging (Vectashield Vibrance Antifade Mounting Medium, Vector Laboratories, H-1700). Images were collected using an FV3000 confocal microscope (Olympus) and further processing was performed using FiJi (Schindelin *et al*, 2012). Confocal image stacks were stitched as two-dimensional maximum intensity projections using FiJi. The mean gray value was measured for the region of interest in the granular cell layer (GCL) of DG and subtract with the image background on each of 2 or 3 slices per brain. Statistical analysis was performed using pairwise Wilcoxon rank sum test.

### Statistical analysis

Fischer’s exact test was used in Figure 2D. The Kruskal–Wallis test was performed for significance in violin plots in Figures S2B and S5C (ns: non-significant; *: *p* < 0.01; **: *p* < 0.001; ***: *p* < 0.0001). Significance in violin plots in Figures 3C, 4C and 5F, and on heatmap in Figure 4B were shown on the basis of adjusted *p*-values obtained with Seurat’s *FindMarkers* function with the MAST algorithm using a hurdle model as a statistical test (ns: non-significant; *: *p* < 0.05; **: *p* < 0.01; ***: *p* < 0.001; ****: *p* < 0.0001). Pairwise Wilcoxon rank sum test was applied to find significant differences in BACH2 expression between young and aged samples.

## Acknowledgements

We would like to thank K. Igarashi and K. Ochiai (Tohoku University) for their advice regarding the analysis for BACH2; The University of Tokyo IQB Olympus Bioimaging Center (TOBIC) for microscopy; E. Ogawara (The University of Tokyo) and V. Panfil (The University of Tokyo and Karolinska Institutet) for technical assistance; and members of Gotoh and Kishi laboratories for helpful discussion. This research was supported by Japan Agency for Medical Research and Development (AMED)-PRIME (JP22gm6110021 to Y.K.), MEXT/JSPS (the Japan Society for the Promotion of Science) KAKENHI (16H06279, 23H04214, 24H01227, and 24K02020 to Y.K.), the Takeda Science Foundation, the Uehara Memorial Foundation, the Asahi Glass Foundation, the Chugai Foundation for Innovative Drug Discovery Science, the Astellas Foundation for Research on Metabolic Disorders, the Naito Foundation, and the SECOM Science and Technology Foundation.

## Author Contributions

**Merve Bilgic**: conceptualization, software, formal analysis, investigation, data curation, writing – original draft preparation, writing – review & editing, visualization, and project administration. **Yukiko Gotoh**: writing – review & editing, and supervision. **Yusuke Kishi**: conceptualization, writing – original draft preparation writing – review & editing, supervision, project administration, and funding acquisition.

## Data and code availability

The sn-multiome-seq dataset was generated from raw sequencing data derived from two replicates of the hippocampus of 7-week-old and 108-week-old mice. Raw datasets (fastq files) have been deposited in the DNA Data Bank of Japan (DDBJ) Sequence Read Archive under the accession code DRAXXX. The external Ogrodnik and Ortiz datasets are available in the GEO database (GSE161340 for Ogrodnik; GSE147747 for Ortiz) (Ogrodnik *et al*, 2021; Ortiz *et al*, 2020). Additional methods and codes used in this study are available from the corresponding author upon reasonable request.

## Declaration of Competing Interests

The authors declare that they have no known competing financial interests or personal relationships that could have appeared to influence the work reported in this paper.

## SUPPLEMENTARY FIGURE LEGENDS

**Figure S1.**
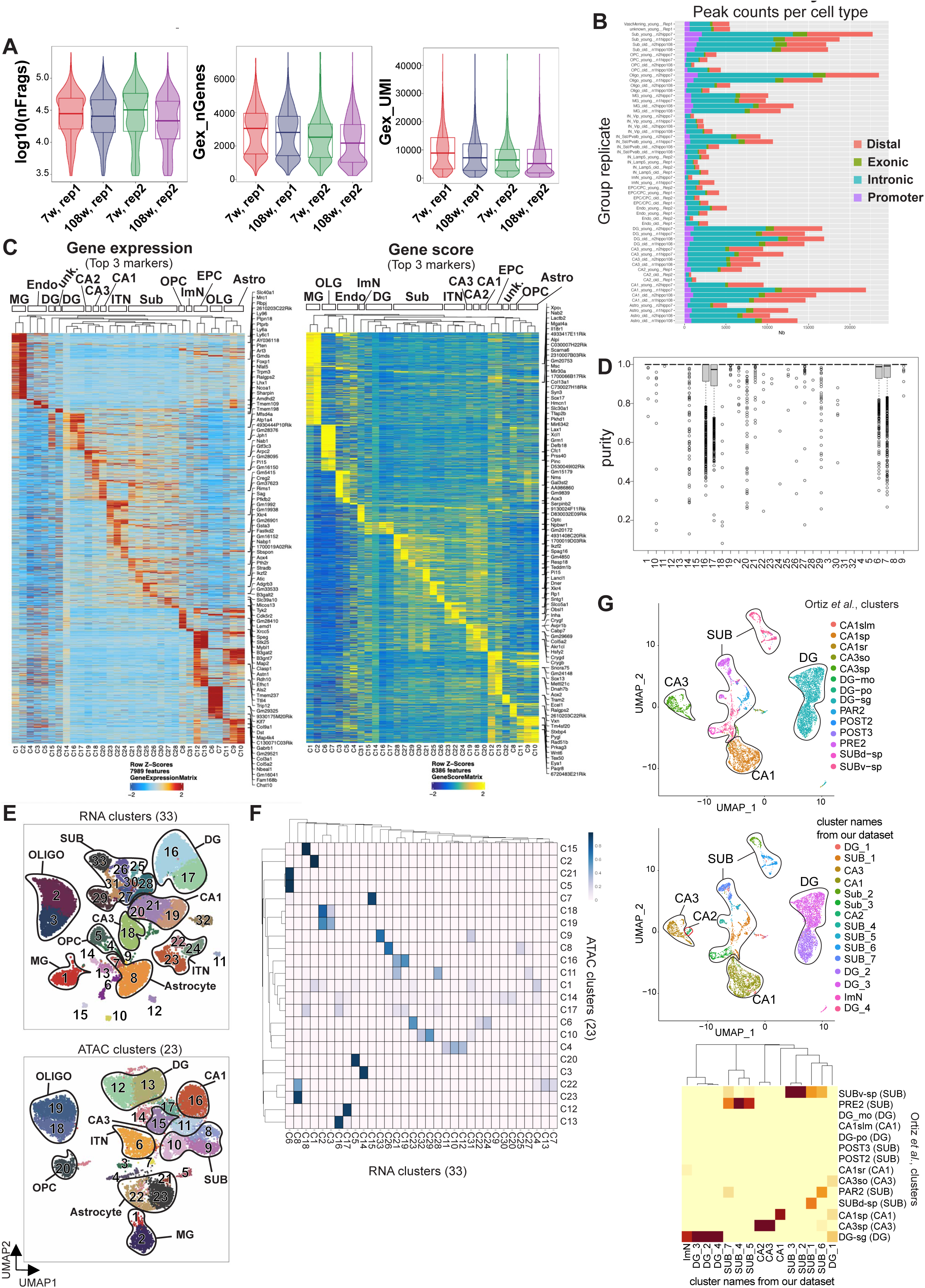
Single-nuclei profiling of transcriptome and chromatin accessibility in the mouse hippocampus with aging. **A.** Quality-control metrics of each replicate in 7-week-old and 108-week-old samples after filtering low-quality cells (Methods). From left to right, violin plots of log10(fragments numbers), gene numbers, and UMI numbers for each cell are shown. **B.** Number of peaks and their annotation is shown in each replicate of cell types from young or aged samples. **C.** Heatmap of top three markers showing gene expression and gene activity in each cluster by hierarchical ordering. Major cell types are indicated in columns above. **D.** Boxplot of cluster purity for each cluster. The purity of neighborhood for each cell was computed and cells are distributed according to their purity along the *y*-axis; the median of each cluster is shown. **E.** Visualization of cells on UMAP colored by RNA or ATAC clusters, with major cell types encircled according to their representative markers. **F.** Heatmap of the confusion matrix representing the distribution of cells across clusters generated by RNA (rows) or ATAC (columns) modalities. Color scale represents the log10(number of cells) for each RNA–ATAC cluster combination. **G.** Visualization of cells colored by predicted cluster identities using Seurat’s label transfer model with the Ortiz et al. dataset, and by cluster identities shown in Figure 1B. The heatmap shows the fraction of predicted identities in ArchR clusters and the color scale indicates the fractions of the range of 0 to 1.

**Figure S2.**
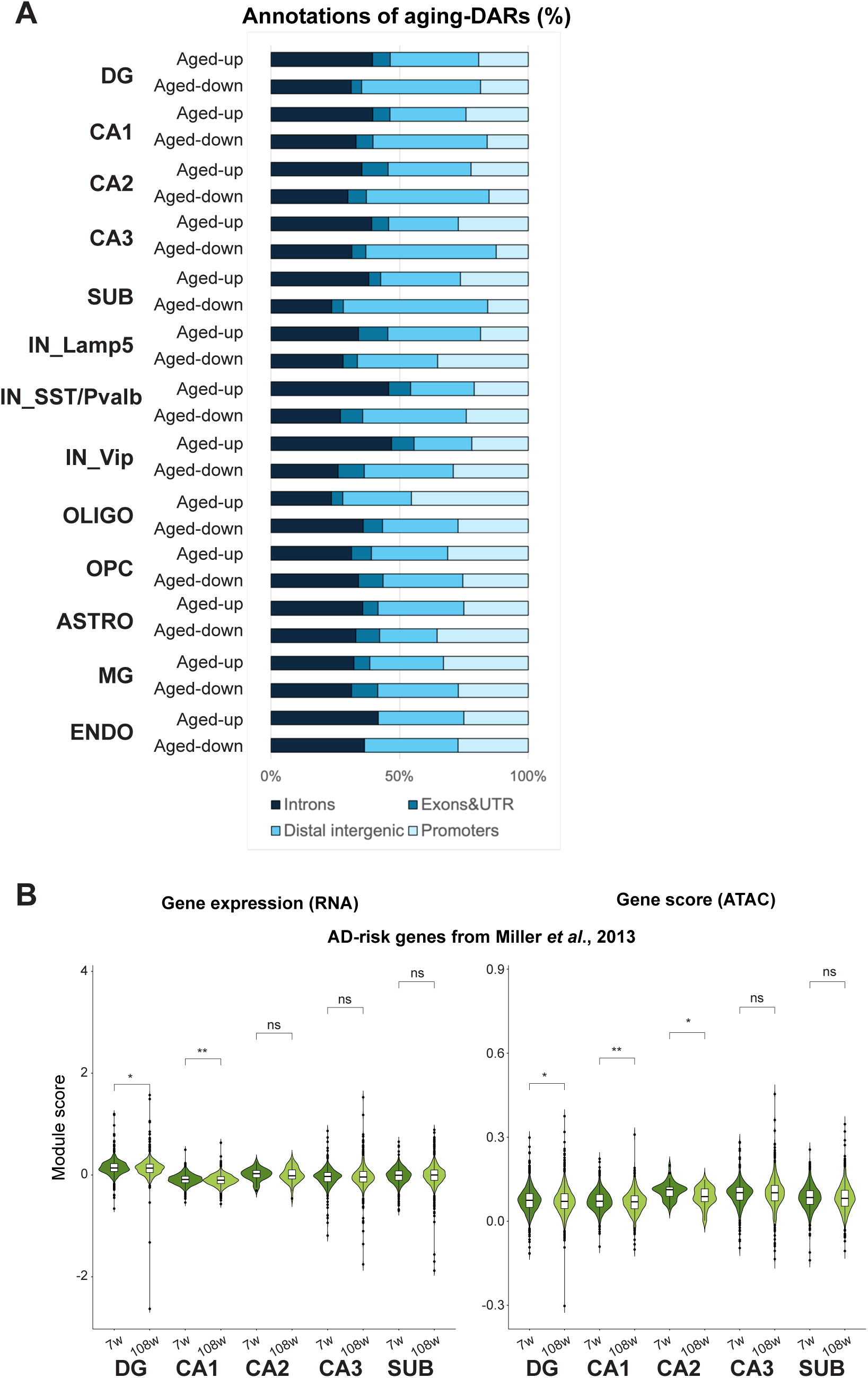
Cell type–specific transcriptome and epigenome dynamics across hippocampal aging. **A.** Percentage of genomic annotations of aging DARs in major cell types. The fractions were calculated separately in aged-up DARs and aged-down DARs. **B.** Module scores for each cell in young and aged DG, CA1-3, and SUB neurons. DEGs identified in CA1 and CA3 of AD hippocampus in Miller et al. were used to calculate module score, representing the average expression levels of these genes in each cell.

**Figure S3.**
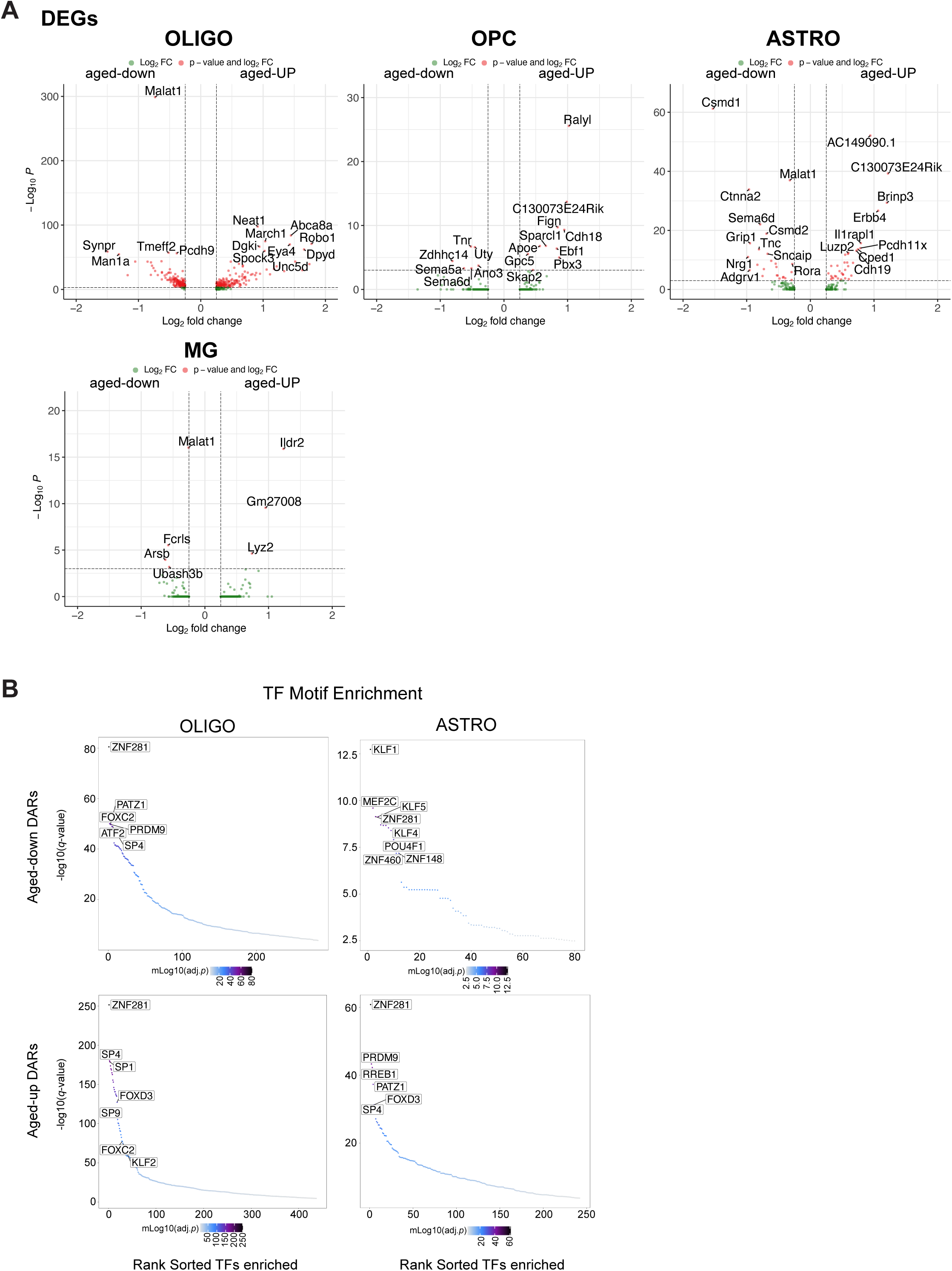
Dysregulation of neuronal genes in glial cells during hippocampal aging. **A.** Volcano plot for DEGs in oligodendrocytes, oligodendrocyte precursors, astrocytes, and microglial cells. The *x*-axis indicates log2(fold-change) of average gene expression, and the *y*-axis indicates significance in −log10(adjusted *p*-value). Vertical threshold is 0.25 and horizontal threshold is 3. **B.** Transcription factor motif enrichment ranked by *E*-values in OLIGO and ASTRO (Methods). TF motif enrichment on aged-down DARs and aged-up DARs are shown separately on upper and lower panels, respectively. The color intensity indicates significance by −log10(adjusted *p*-value).

**Figure S4.**
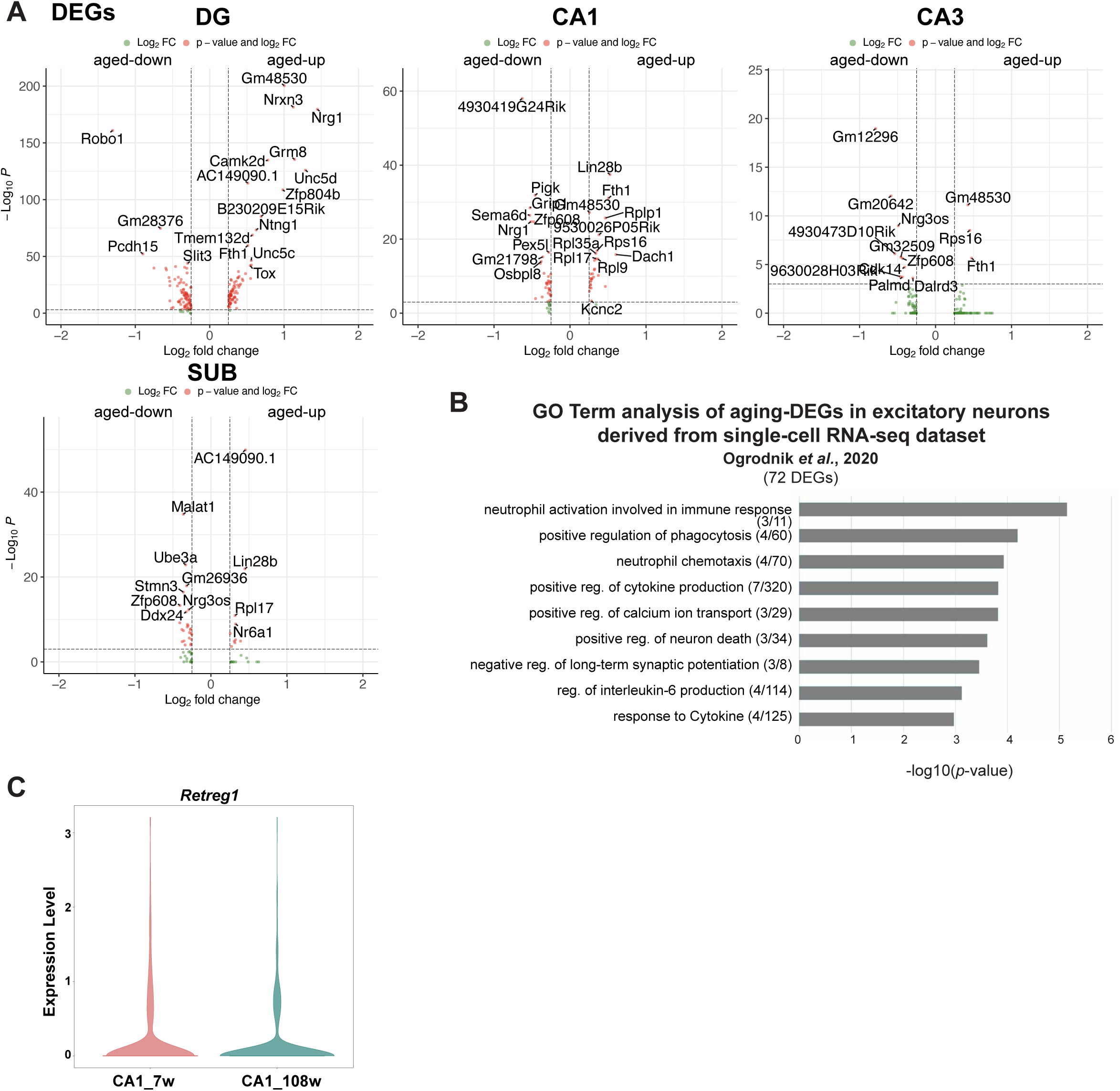
Chromatin accessibility–level dysregulations recapitulated aging features in neurons. **A.** Volcano plot for DEGs in neuronal subtypes including DG, CA1, CA3, and SUB. The *x*-axis indicates log2(fold-change) of average gene expression, and the *y*-axis indicates significance in −log10(adjusted *p*-value). Vertical threshold is 0.25 and horizontal threshold is 3. **B.** Gene ontology enrichment analysis for biological processes using DEGs identified in Ogrodnik single-cell datasets for the aging hippocampus. We identified 72 significant DEGs in excitatory neurons. The *x*-axis indicates the significance by −log10(*p*-value). **C.** Violin plot of gene expression for *Retreg1* in CA1 neurons split by age.

**Figure S5.**
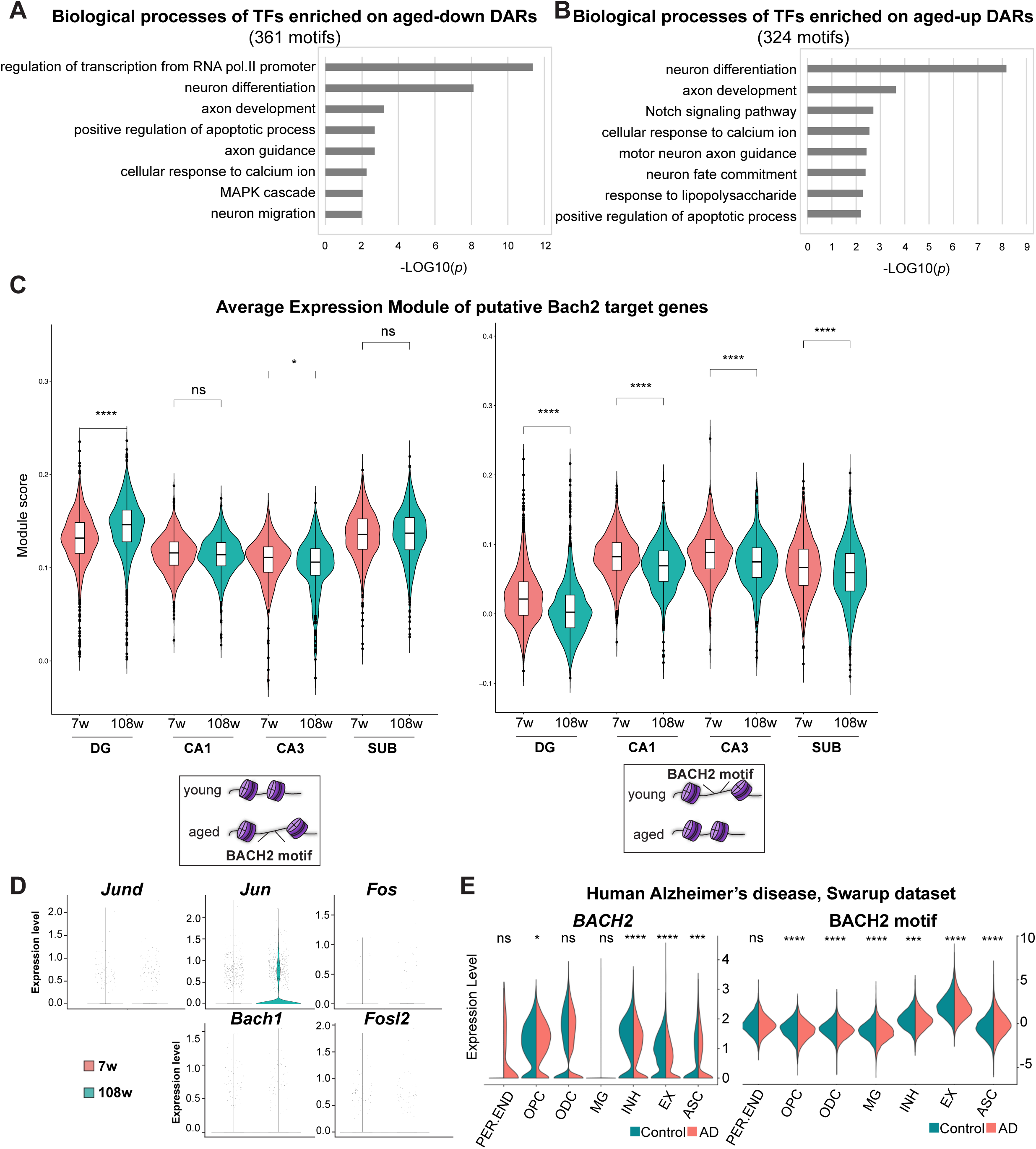
BACH2 activity reduced in aged DG neurons. **A–B**. Gene ontology enrichment analysis for biological processes in neuronal subtypes of TF motifs enriched on aged-downregulated DARs (A) and aged-upregulated DARs (B). **C**. Violin plot of average expression scores for BACH2-putative target genes identified in Figure 5, in each cell of neuronal subtypes split by their age. **D**. Violin plot of average gene expression for *Bach1* and AP-1 components, *Jund*, *Jun*, *Fos*, and *Fosl2* in DG neurons split by their age of collection. **E**. Violin plot of expression level (left) and motif enrichment (right) of Bach2 in human AD dataset. The plots were taken from https://swaruplab.bio.uci.edu/singlenucleiAD/ (Morabito *et al*, 2021).

## Supplementary Data

Supplementary Table 1. Quality control, metadata, and cluster information, related to Figure 1.

Supplementary Table 2. DEG and DAR analyses related to Figure 2.

Supplementary Table 3. TF motif enrichment related to Figure 3.

Supplementary Table 4. Gene Ontology analysis related to Figure 4.

Supplementary Table 5. TF and GO analyses related to Figure 5.

## Notes

### Competing Interest Statement

The authors have declared no competing interest.

